# Antidepressant-like effect of losartan involves TRKB transactivation from angiotensin receptor type 2 (AGTR2) and recruitment of FYN

**DOI:** 10.1101/168708

**Authors:** Cassiano R.A.F Diniz, Plinio C. Casarotto, Senem M. Fred, Caroline Biojone, Eero Castrén, Sâmia R. L. Joca

## Abstract

Renin-angiotensin system (RAS) is associated to peripheral fluid homeostasis and cardiovascular function, but recent evidence has also drawn its functional role in the brain. RAS has been described to regulate physiological and behavioral parameters related to stress response, including depressive symptoms. Apparently, RAS can modulate levels of brain derived neurotrophic factor (BDNF) and TRKB, which are important to neurobiology of depression and antidepressant action. However, interaction between BDNF/TRKB system and RAS in models predictive of antidepressant effect has not been investigated before. Accordingly, in the forced swimming test, we observed an antidepressant-like effect of systemic losartan but not with captopril or enalapril treatment. Moreover, infusion of losartan into ventral hippocampus (vHC) and prelimbic prefrontal cortex (PL) mimicked the consequences of systemically injected losartan, whereas K252a, a blocker of TRK, infused into these brain areas impaired such effect. PD123319, an antagonist of AT2 receptor (AGTR2), infused into PL but not into vHC, also prevented systemic losartan effect. Cultured cortical cells of rat embryos indicate that angiotensin II (ANG2), possibly through AGTR2, increases the surface levels of TRKB, and favors it’s coupling to FYN, a SRC family kinase. The higher levels of *agtr2* in cortical cells were decreased after insult with glutamate, and under this condition an interaction between losartan and ANG2 was achieved. Occurrence of TRKB/AGTR2 heterodimers was also observed, in MG87 cells GFP-tagged AGTR2 co-immunoprecipitated with TRKB. Therefore, antidepressant-like effect of losartan is proposed to occur through a shift of ANG2 binding towards AGTR2, followed by coupling of TRK/FYN and putative TRKB transactivation. Thus, AGTR1 show therapeutic potential as novel antidepressant therapy.

## INTRODUCTION

Renin-angiotensin system (RAS) functional role has been historically implicated in cardiovascular and fluid homeostasis. Firstly, precursor molecule angiotensinogen is cleaved by renin into angiotensin I, which is then converted into Angiotensin II (ANG2) by angiotensin-converting enzyme (ACE) (Wright and Harding, 2011). Main actions of the ANG2 are mediated by angiotensin II receptors type 1 and 2 - AGTR1 and AGTR2, respectively (Wright and Harding, 2011).

Other reports, however, has pointed out all components of renin-angiotensin being produced inside central nervous system (CNS) (Saavedra, 2005). Thus, AGTR1 and AGTR2 in circumventricular organs and in cerebrovascular endothelial cells may respond to circulating ANG2 of peripheral origin, whereas receptor lying in neurons inside blood brain barrier respond to RAS of brain origin (Saavedra, 2005). AGTR1 and AGTR2 have been found expressed inside blood brain barrier structures such as hippocampus and frontal cortex (Bunnemann et al., 1992; Tsutsumi and Saavedra, 1991), both considered crucial limbic structures associated to the neurobiology of depression (Krishnan and Nestler, 2008).

In fact, several piece of evidences introduce ANG2 as a hormone regulator of peripheral and central physiological changes regarding stress exposure, including behavioral consequences. For instance, both acute and chronic stress increased ANG2 and AGTR1 expression levels in the hypothalamic-pituitary-adrenal axis (HPA axis) (Castren and Saavedra, 1988; Saavedra et al., 2004; Yang et al., 1993). Moreover, candesartan (AGTR1 antagonist) treatment prevented stress effect of increasing pituitary adrenocorticotrophic and adrenal corticosterone hormone levels (Armando, 2001), and treatment with ACE inhibitors (ACEi) or AGTR1 antagonists reversed or prevented animal behavioral responses to stress (Gard et al., 1999; Giardina and Ebert, 1989; Martin et al., 1990a, 1990b; Ping et al., 2014; Vijayapandi and Nagappa, 2005). In the same way, animals lacking angiotensinogen showed antidepressant-like phenotype (Okuyama et al., 1999).

The neurotrophin brain-derived neurotrophic factor (BDNF), found mostly in the central nervous system, is important for neural plasticity, including synapse formation, neuronal differentiation and growth (Park and Poo, 2013). Functional role of BDNF and its receptor (TRKB, *tropomyosin-related kinase B receptor*) has been linked with pathophysiology of psychiatric disorders, such as depression, and with mechanism of action of antidepressant drugs (Castrén, 2014). Apparently, RAS may modulate BDNF and TRKB brain levels. For instance, candesartan treatment prevented both infarct volume and neurological deficit in animals suffering middle cerebral artery occlusion while increased protein and mRNA levels of TRKB in the brain (Krikov et al., 2008). In addition, telmisartan (AGTR1 antagonist) chronic treatment was able to prevent retinal damage and decrease of BDNF levels, such as observed in diabetic animal model (Ola et al., 2013). Valsartan, another AGTR1 antagonist, counteracted the consequences of stress on depressive and anxiogenic-like behavior, as well as on BDNF levels in hippocampus and frontal cortex (Ping et al., 2014). Moreover, some case reports describe relief of depressive symptoms in hypertensive patients treated with the ACEi captopril (Deicken, 1986; Germain and Chouinard, 1988; Zubenko and Nixon, 1984).

Despite scarce evidence, it is plausible to consider that drugs acting on RAS promote antidepressant-like effects. However, such properties have not been linked to the modulation of BDNF/TRKB system. In this sense, the present work aimed at investigating behavioral effects of AGTR1 antagonist losartan and ACEi in a model predictive of antidepressant-like effect, i.e. forced swimming test, and the requirement of BDNF/TRKB for such effect. Since AGTR1 activation was related to brain injury (Saavedra, 2012) and activation of AGTR2 has been supposed to employ balancing neuroprotective outcomes, especially when AGTR1 are blocked (Mogi and Horiuchi, 2013; Zhao et al., 2005), we hypothesized that activation of AGTR2 could underlie the antidepressant-like effects of losartan. In vitro analysis from cultured cortical cells were also performed to provide a mechanistic insight to the behavioral data.

## MATERIAL AND METHODS

### Animals

Male Wistar rats (250-350g), and female C57BL6/j background mice, heterozygous to BDNF expression or WT littermates were used in behavioral studies. The rats were housed in pairs while mice were group housed (4-6/cage) in a temperature controlled room (24±1°C) under standard laboratory conditions with access to food and water *ad libitum* and a 12h light/12h dark cycle (light on at 6:30a.m.). *In vivo* experiments were conducted in conformity with local Ethical Committee (protocol 147/2017 for USP and ESAVI/10300/04.10.07/2016 for UH), which is in accordance with Brazilian Council for the Control of Animals under Experiment (CONCEA), European Union and ARRIVE guidelines (Kilkenny et al., 2010)for the care and use of laboratory animals. All comply with international laws and policies.

### Cell Culture

Mouse fibroblasts stably overexpressing full-length TRKB (MG87.TRKB) were cultured in Dulbecco’s Modified Eagle’s Medium (DMEM) (supplemented with 10% fetal calf serum, 1% penicillin/streptomycin, 1% l-glutamine and 400 mg/ml G418). Cell lines were maintained at 5% CO2, 37°C until reaching 70% confluence for experiments. For primary neuronal cultures, cortices from E18 rat embryos were dissected and had tissue dissociated with papain solution in PBS (10min, 37°C). Cells were suspended in DMEM medium containing Ca^2+^/Mg^2+^ free HBBS, 1mM sodium pyruvate, 10mM HEPES (pH 7.2) and DNAse, and plated onto poly-l-lysine (Sigma–Aldrich) coated 24- or 96-well culture plates at a cell density of 125000 cells/cm^2^. Primary neurons were maintained in Neurobasal medium (supplemented with 2% B27, 1% penicillin/streptomycin and 1% l-glutamine) and supplemented with fresh medium every 3rd day.

### Drug Treatments

Losartan potassium (losartan; AGTR1 antagonist; Pharmanostra, BRAZIL) was administered intraperitoneally - ip (10, 30 and 45mg/Kg), intracerebrally (0.1, 1 and 10nmoL/site) or in cell cultures (10uM; Tocris #3798). Captopril (ACEi; 3, 10 and 30mg/Kg - Pharmanostra, BRAZIL) and enalapril (ACEi; 1, 3 and 10mg/Kg - Pharmanostra, BRAZIL) were injected ip. K252a was used for intracerebral (TRK inhibitor; 20mM/site – Sigma-Aldrich, #K2015) or in culture (10uM) treatment. PD123319 (PD; AGTR2 antagonist; Tocris, #1361) was used for intracerebral (200uM/site) or culture (200nM) treatment. ANG2 (10uM; Tocris #1158), BDNF (2.5ng/mL), NGF (10ng/mL), TRKB.Fc (200ng/mL; R&D systems, #688-TK-100) and CGP42112 (CGP 2uM, AGTR2 agonist, Tocris, #2569) were used only for cell culture treatment. K252a and PD123319 intracerebral doses were chosen based on previous works (Jiang et al, 2012; Zheng et al, 2008). 2,5% 2,2,2 tribromoethanol (ip, Sigma-Aldrich, #T48402) and subcutaneous local anesthetic lidocaine (PROBEM 3%, 0.2 mL) were used for stereotaxic surgery. Chloral hydrate (0.75g/Kg, ip, Sigma-Aldrich, #C8383) was used to euthanize animals for perfusion. Subcutaneous banamine (Schering-Plough, 0.25%, 0.1mL/100g) and intramuscular oxytetracycline (Pfizer, 20%, 0.1mL/100g) were used once to postoperative recovery. Losartan, tribromoethanol, choral and banamine were freshly prepared in in saline solution, whereas all other drugs in 0.1% DMSO in saline.

### Surgery, Intracerebral Injections and Histology

Surgery and intracerebral drug injections were performed as described earlier (Diniz et al., 2016). Briefly, rats were anesthetized with tribromoethanol and fixed in a stereotaxic frame. Further, stainless steel guide cannulas (0.7mm OD) aimed at the dorsal hippocampus (dHC; coordinates: AP= −4.0mm from bregma, L=2.8mm, DV=2.1mm), ventral hippocampus (vHC; coordinates: AP= −5.0mm from bregma, L=5.2mm, DV=4.0mm) or prelimbic ventromedial prefrontal cortex (PL; coordinates: AP= +3.3mm from bregma, L=1.9mm, DV=2.4mm; lateral inclination of 22°) were implanted according to Paxinos and Watson’s atlas (Paxinos and Watson, 1998) and attached to skull bone with stainless steel screws and acrylic cement. A stylet inside guide cannula prevented obstruction. Five to seven days after surgery, intra-cerebral injections were performed with dental needle (0.3mm OD) in a volume of 200nl (mPFC) or 500nl (dHC or vHC) infused for 1min using a Hamilton micro-syringe (Sigma-Aldrich) and infusion pump (KD Scientific). After behavioral tests, rats were anesthetized with chloral hydrate and 200nL of methylene blue was injected through the guide cannula. The brains were removed and injection sites verified. Results from injections outside the aimed area were discarded from statistical analysis. All histological sites of injection were inserted in diagrams (Figure S2) based on the atlas of Paxinos and Watson (Paxinos and Watson, 1998).

### Forced Swimming Test (FST)

The FST was performed in rats as follows: animals were placed individually to swim in a Plexiglas cylinder (24cm diameter by 60cm with 28cm of water at 25±1°C) for 15min (pretest). Twenty-four hours later, animals were replaced in the cylinder for 5min swim test session and immobility time was measured (Diniz et al., 2016).

The FST in mice was performed as follows: the animals were placed in 5l glass beaker cylinders (19cm diameter, with 20cm water column for 6min. The immobility was assessed in the last 4min of the session (Diniz et al., 2017). For both protocols, water was changed between each test. After swimming, animals were towel-dried and kept in a warmed cage before returning to home cages. Test was videotaped and analyzed by a trained observer blind to treatment.

### Overexpression of GFP-AGTR2

MG87 cells were transfected to express GFP-tagged AGTR2 using lipofectamine (Jiang et al, 2012). Briefly, at a confluence of 70%, the cells were incubated with a mixture of 2.5% lipofectamine 2000 (Thermo Scientific, #11668019) and 5ug/mL of the plasmid in OptiMEM medium. Following 48h after transfection, cells were treated, lysed and submitted for immunoprecipitation as described below.

### Sample collection

For immunoassays, cells were washed with ice-cold PBS, lysed [137mM NaCl; 20mM Tris-HCl; 1%NP40; 0.5mM NaF; 10% glycerol; pH=7.4; supplemented with protease/phosphatase inhibitor cocktail (Sigma-Aldrich, #P2714; #P0044) and sodium orthovanadate (0.05%, Sigma-Aldrich, #S6508] and the samples were centrifuged at 10000g for 15min at 4°C. Supernatant was collected and stored at −80°C until use. For polymerase chain reaction, the cells were washed with PBS and treated with Qiazol Lysis Reagent^™^ (Qiagen). Lysate was collected in a clean tube and incubated with chloroform for 3min at RT. After centrifuged at 15200g for 10min at 4°C, the aqueous phase was mixed with isopropanol for 10min at RT and centrifuged at 15200g for 10min at 4°C. The pellet was washed with 75% EtOH two times, than with 100% EtOH once, air-dried and dissolved in 20μL MQ-water.

### Reverse Transcription-Polymerase Chain Reaction (RT-PCR) and Quantitative Polymerase Chain Reaction (qPCR)

Concentration and purity of each RNA sample were determined using NanoDrop (Thermo Scientific). Maxima First Strand cDNA Synthesis Kit for RT-qPCR with dsDNase (#K1672, Thermo Scientific) was used to synthesize cDNA from the samples. Primers were designed via https://eu.idtdna.com/pages/scitools and purchased from Sigma-Aldrich (Germany). The following primers were used for qPCR:

AGTR1a-(NM_030985.4) forward: CATCAGTCTCCCTTTGCTATGT, reverse: AGTGACCTTGATTTCCATCTCTT.

AGTR2-(NM_012494.3) forward: CCTTCCATGTTCTGACCTTCTT, reverse: GCCAGGTCAATGACTGCTATAA.

Actin-(NM_031144) forward: TGTCACCAACTGGGACGATA, reverse: GGGGTGTTGAAGGTCTCAAA.

The PCR method used SYBR Green as probe. Briefly, maxima®SYBR Green qPCR Master Mix (Thermo Scientific, #K0253) was used according to manufacturer’s instructions in Hard-Shell^™^ 96-well PCR plate (BioRad). The reaction was conducted in duplicates, using thermal cycler (BioRad CFX96 Real-Time System), with initial denaturation at 95°C for 10min. Denaturation and amplification were carried out by 45 cycles of 95°C for 15s, 63°C for 30s and 72°C for 30s. ‘No template control’ (NTC) was included to the reaction and melting curve analysis was done. Results were analyzed using Ct values. Levels of beta-actin mRNA was used for normalization of the results and 2^−ΔΔCt^ values were calculated for each gene.

### Immunoprecipitation

Lysate from transfected MG87.TRKB/GFP-AGTR2 cells were incubated with antibody against TRK (Santa Cruz, #sc7268) overnight (1ug of Ab: 500ug of total proteins) at 4°C. Following incubation with Protein-G Sepharose (Life Technologies, #101242) for 2h at 4°C, samples were centrifuged (10000g/2min) and the precipitate was stored at −80°C until use. This cycle was repeated twice, using lysis buffer to wash samples. Supernatant was discarded.

### Western-blotting and ELISA

For western-blotting, protein precipitated by anti-TRK and supernatant were separated in SDS-PAGE and transferred to PVDF membranes. Following blocking with 3% BSA in TBST, membranes were incubated with antibody against GFP (1:1000, Santa Cruz, #sc-8334) or total TRK (1:1000, Santa Cruz, #sc11-Rb). Membranes were incubated with secondary antibody conjugated to HRP (1:10000, Bio-Rad, #170-5046) and chemiluminescence emitted after addition of ECL was detected by CCD camera. Immunoblot bands were measured using NIH ImageJ 1.32.

For ELISA, samples (120ug total proteins) were incubated (overnight at 4°C) in 96-well plates, previously coated with anti-TRK (1:500, Santa Cruz, #sc7268, overnight at 4°C) and blocked with 3%BSA (2h at RT) in PBST. Following wash with PBST, anti-pTRK.Y816 (1:2000, Cell Signaling, #4168), biotin-conjugated anti-pY (1:2000, AbD Serotec, UK, #MCA2472B) or anti-FYN (1:2000, Santa Cruz, #sc16) was incubated overnight at 4°C. After wash with PBST, plate was incubated with HRP-conjugated tertiary antibody (1:5000, Bio-Rad, #170-5046) or HRP-conjugated streptavidin (1:10000, Thermo Fisher, #21126). Chemiluminescence emitted after addition of ECL was detected by a plate reader (Varioskan Flash, Thermo-Fisher). Signal from each sample, discounted blank, was normalized and expressed as percentage of control-group.

### Surface expression of TRK

Cells from rat E18 cortex were cultivated in 96-well plates as described above (DIV8). Detection of surface TRKB was performed by ELISA (Zheng et al., 2008). Cells were fixed with 4%PFA for 20min at RT. After washing with PBS wells were blocked with 5% non-fat dry milk and 5% normal goat serum in PBS for 1h at RT. Then, primary antibody against extracellular portion of TRK (1:500, Santa Cruz, #sc8316) was incubated overnight at 4°C. Following wash with PBS, cells were incubated with HRP-conjugated antibody (1:5000; Bio-Rad, #170-5046) for 2h at RT. Signal detected after addition of ECL, discounted blank, was normalized by the average of vehicle-treated samples and expressed as percentage of control-group.

### Data Analysis

Statistical analyses were carried out using two-tailed Student’s t-test, one-way analysis of variance (ANOVA) followed by Fisher’s LSD *post-hoc* test, or two-way ANOVA test. Criteria for statistical significance was p<0.05.

## RESULTS

### Antidepressant-like Effect of Losartan

Different classes of drugs, with discrepant mechanisms concerning RAS modulation, were used to evaluate a possible drug-induced antidepressant effect. As depicted in figure 1, one-way ANOVA indicates a significant effect of treatment losartan (F_4,46_=8.27, p<0.05; Figure 1c) both losartan at 10 and 45mg/kg dose and imipramine (30mg/kg) reduced the immobility time in FST (Fisher’s LSD p<0.05 for all).. Both captopril (F_3,29_=1.83, non-significant NS; Figure 1a) or enalapril (F_3,16_=0.46, NS; Figure 1b) treatment were not effective in decreasing the immobility time in the FST. In fact, immobility time of rats exposed to swimming session is increased after uncontrollable stress and the treatment with antidepressant drugs decrease this parameter (Nestler and Hyman, 2010). Moreover, provided that known antidepressants decrease immobility time in FST, a good predictive validity is attributed to it, thus supporting FST as a screening test for putative new antidepressant drugs and their mechanisms. Since only systemic losartan exhibited antidepressant-like effect, this drug was infused into dHC, vHC or PL to address which of these structures may underlie such effect. The data depicted in figure 1d-f indicates that losartan infused into vHC (F_3,11_=10.66, p<0.05; Figure 1e) and PL (F_3,19_=2.72, p<0.05; Figure 1f), but not into dHC (F_3,24_=0.50, NS; Figure 1d), was able to decrease immobility time in the FST.

**Figure 1.**
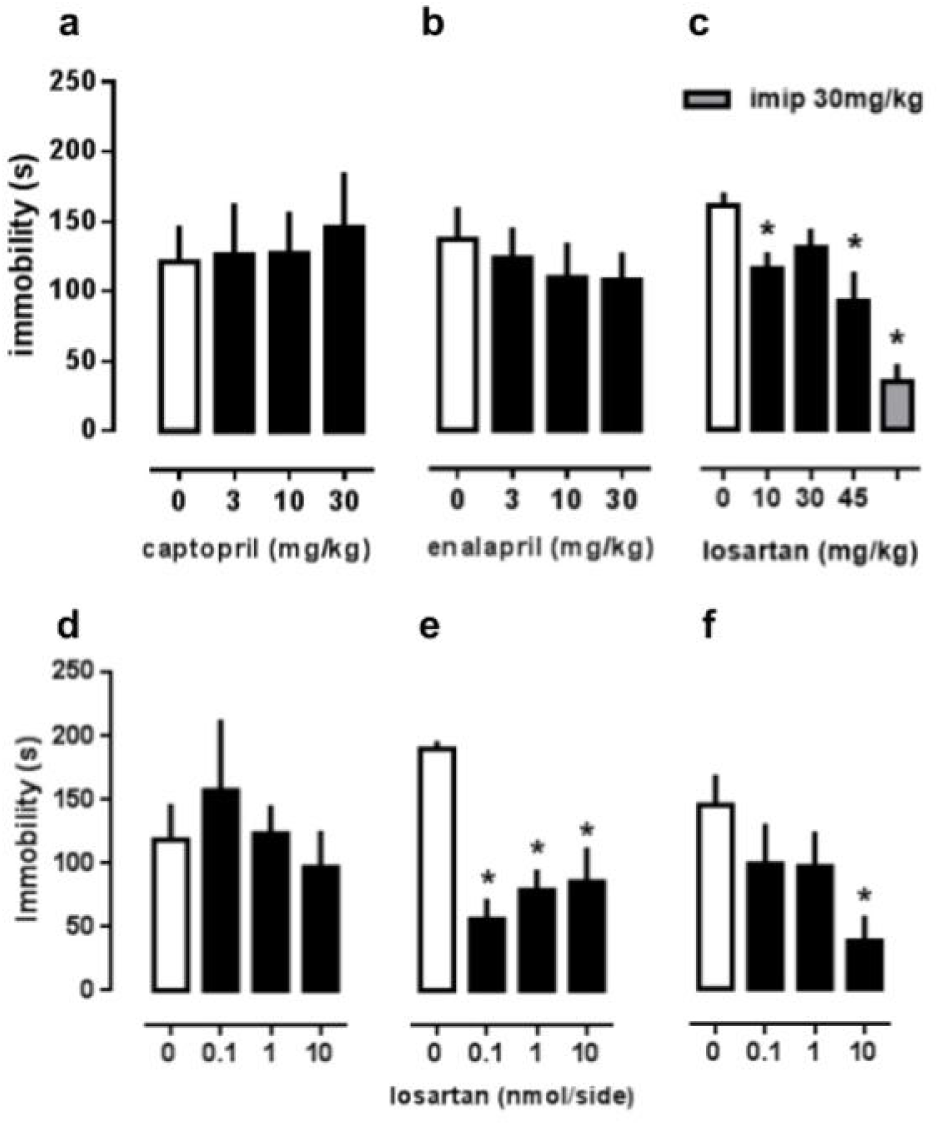
Antidepressant-like Effect of Losartan. Animals were treated i.p 24h, 5h and 1 hour before FST with captopril (a) at 0, 3, 10 or 30mg/kg (n=7-9/group) doses or with enalapril (b) at 0, 1, 3 or 10mg/kg (n=5/group) doses. (c) Losartan was administered i.p 1h before FST at 0, 10, 30 or 45mg/kg and imipramine at 30mg/kg was used as positive control (n=5-16/group). (d-f) Losartan was bilaterally infused at 0, 0.1, 1 and 10nmol/side into (d) dorsal hippocampus - dHC (n=4-8/group), (e) ventral hippocampus - vHC (n=3-4/group) or (f) prelimbic prefrontal cortex - PL (n=5-7/group). Data are expressed as mean ± SEM of immobility time (s); *p<0.05 compared to control group.

### Interaction between losartan and TRK or AGTR2: *in vivo* data

In this experimental set, requirement of TRK and/or AGTR2 activation for antidepressant effect of losartan was examined. As shown in figure 2a-b, K252a, an antagonist of TRK receptors, infused into vHC or PL was able to modulate the effect of systemically injected losartan. Two-way ANOVA revealed interaction between factors (systemic and intracerebral injection) for both structures (vHC: F_1,15_=8.62; PL: F_1,13_=6.58, p<0.05 for both). Regarding vHC experiments, pairwise comparisons unveiled immobility time of animals treated with losartan/ctrl is decreased compared to control group (Fisher’s LSD, p<0.05), whereas losartan/K252a group was considered significantly different of losartan/ctrl group (Fisher’s LSD, p<0.05), suggesting that K252a prevents antidepressant-like effect of losartan. Regarding PL experiments, pairwise comparisons unveiled immobility time of animals treated with losartan/ctrl is lower than control group (Fisher’s LSD, p<0.05), whereas losartan/K252a group was found different from losartan/ctrl group (Fisher’s LSD, p<0.05), suggesting that K252a also prevents losartan effect in this brain region.

**Figure 2.**
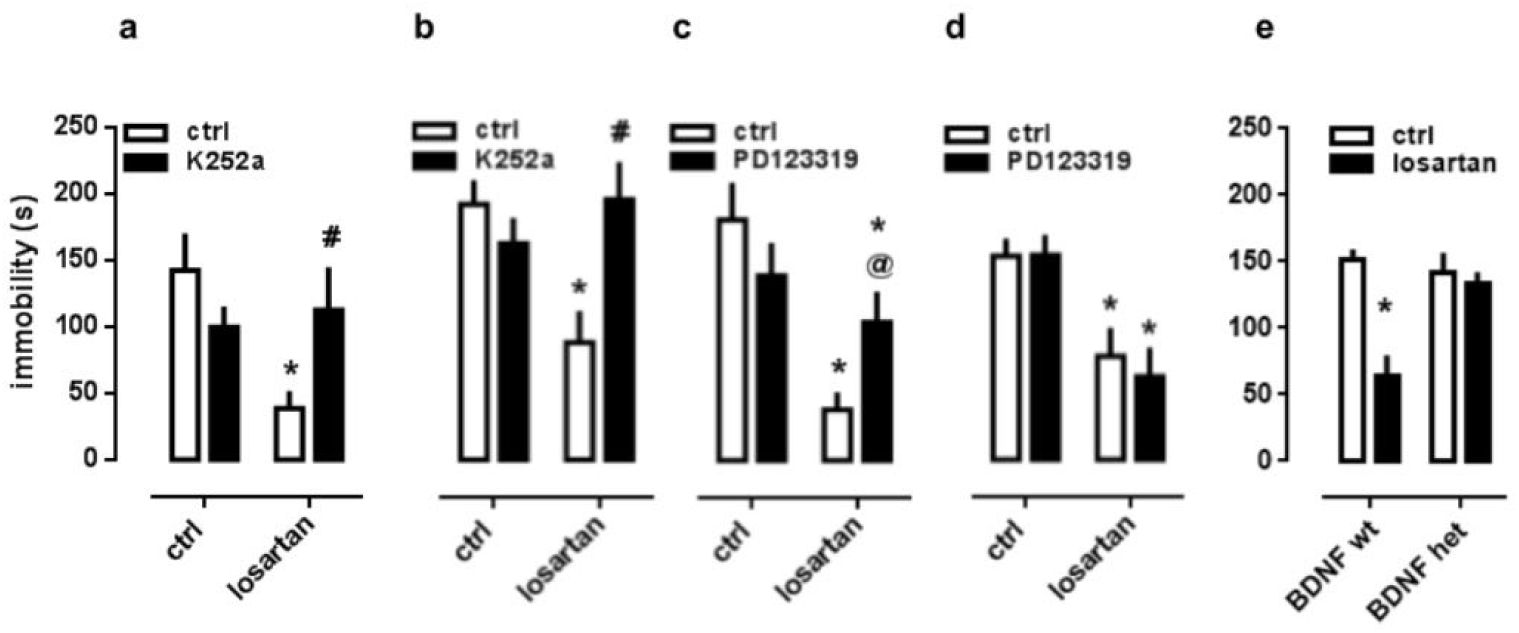
Interaction between losartan and TRK or AGTR2: *in vivo* data. Antidepressant-like effect of losartan was attenuated by K252a infused into (a) PL (n=4-5/group) and (b) vHC (n=4-6/group). The infusion of AGTR2 antagonist PD123319 into (c) PL (n=7-9/group) attenuated the effect of systemically injected losartan, but no change was observed in (d) vHC (n=4-6/group). (e) Interaction between genotype and losartan effect in mice haploinsufficient to BDNF or WT littermates (n= 5-8/group). Losartan was administered 1h before FST and either K252a or PD123319 was bilaterally infused 20min before swimming test. Data are expressed as mean ± SEM of immobility time (s). *p<0.05 compared to ctrl/ctrl group, unless otherwise stated; #p<0.05 compared to PD/VEH group. @p=0.06 compared to PD/VEH group.

Next, we analyzed if antidepressant effect of losartan relies on AGTR2 activity. As shown in figure 2c-d, the AGTR2 antagonist PD123319 infused into PL, but not into vHC, was able to mitigate the effect of systemically injected losartan. Accordingly, Two-way ANOVA revealed interactions between the compounds in PL (F_1,28_=5.11, p<0.05) but not in vHC (F_1,26_=0.18, NS) structure. In addition, pairwise comparisons concerning PL experiments unveiled immobility time of animals treated with losartan/ctrl is reduced compared to control group (Fisher’s LSD, p<0.05), however losartan/PD123319 group was not found different from losartan/ctrl (Fisher’s LSD: t_28_=1.95, p=0.06). Therefore, activation of AGTR2 in PL, but not in vHC, is necessary to mediate the antidepressant-like effect of losartan.

Finally, we submitted BDNF haploinsufficient mice and their littermate (WT) controls to FST. As shown in fig X. The administration of losartan decreased the immobility time only in WT animals, while no effect was observed in animals with reduced BDNF levels [interaction: F_1,24_= 10.15; p<0.05], figure 2e. Considering only the WT group, the one-way ANOVA indicates that both losartan (45mg/kg) and fluoxetine (30mg/kg; used as a positive control) reduced the immobility in the FST [mean immobility time (s)/SEM(n). ctrl: 150.6/8.39(7); losartan: 64.25/15.40(8); fluoxetine: 39.8/9.65(5). F_2,17_=20.44; p<0.05]; data not plotted.

### Interaction between TRK and AGTR2: *in vitro* data

Since losartan antidepressant-like effect may depend mutually on TRKB and AGTR2 signaling in PL, we hypothesized losartan treatment would allow a shift of ANG2 from AGTR1 towards AGTR2 to forward TRKB activation. In order to test this possibility, losartan, PD123319 or K252a (1^st^ factors) was added to the primary cell cultures previously to ANG2 (2^nd^ factor) and levels of pTRK were analyzed. In fact, two way ANOVA indicated an interaction between both factors (F_3,28_=6.77, p<0.05) and pairwise comparisons assert that both PD123319 and K252a, prevented ANG2 effect of increasing pTRK levels (Fisher’s LSD, NS for both; Figure 3a) and no additive effect was observed in the combination of ANG2 and losartan. Next step was to check if ANG2 effect was dependent on BDNF release. With this intent, a soluble BDNF scavenger - TRKB.Fc (1^st^ factor) - was added to the medium of the cell culture before ANG2 (2^nd^ factor). Two way ANOVA indicated no interaction between both factors (F_1,20_=0.82, NS), suggesting an effect of ANG2 on pTRK levels is independent of BDNF release (Figure 3b).

**Figure 3.**
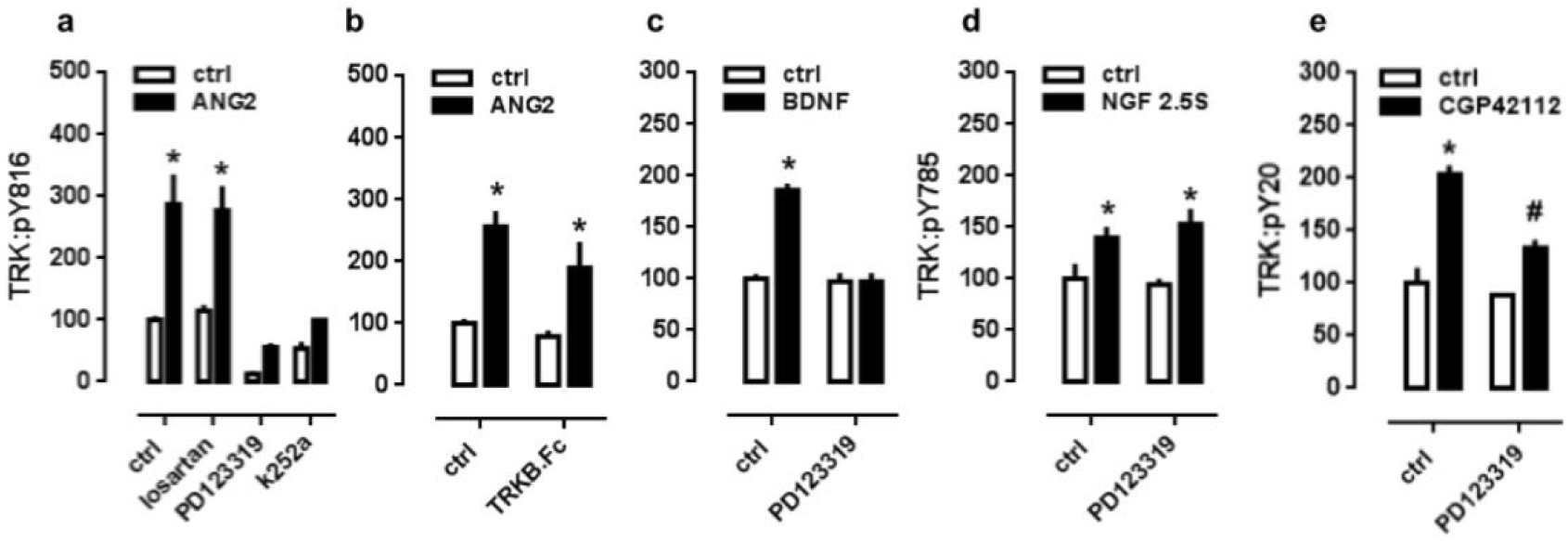
Interaction between TRK and AGTR2: *in vitro* data. (a) Previous administration of PD or K252a, but not losartan, blocked the ANG2-induced increase in TRKB activation in cortical cells of rat embryo - E18; DIV8-10 (n=4-6/group). (b) previous incubation with TRKB.Fc did not change ANG2-induced increase in TRKB activation in cortical cells (n=6/group). Previous incubation with PD123319 blocked (c) BDNF- and (e) CGP42112-, but not (d) NGF-induced activation of TRKB (n=4-6/group). *p<0.05 compared to ctrl/ctrl group; cells were lysed 10min after last drug administration.

To further investigate that, PD123319 (1^st^ factor) was added to culture medium before BDNF (2^nd^ factor) and levels of pTRK were analyzed. Two-way ANOVA showed a significant interaction between factors (F_1,20_=36.18, p<0.05). Pairwise comparisons indicate PD123319 abrogated BDNF effect of increasing pTRK levels (Fisher’s LSD, p<0.05 p<0.05; Figure 3c), indicating AGTR2 participates in BDNF-induced TRK activation. In addition, previous administration of PD123319 (1^st^ factor) was not able to prevent NGF (2^nd^ factor) effect of increasing pTRK levels (Figure 3d), since no interaction between both factors was observed (F_1,13_=0.60, NS). The administration of AGTR2 agonist CGP 42112 was also able to increase the levels of pTRK, and such effect was blocked by previous administration of PD123319 [interaction: F_1,20_= 9.78; p<0.05], figure 3e.

### AGTR2-dependent interaction with FYN, surface positioning of TRKB and co-immunoprecipitation of AGTR2 and TRKB

Inasmuch as ANG2 increased pTRK levels independent of BDNF release, we decided to verify if ANG2 and BDNF are able to influence TRK/FYN coupling, since FYN is described as a SRC member responsible for transactivation of TRK (Rajagopal and Chao, 2006). For this purpose, PD123319 (1^st^ factor) was added in the culture medium before ANG2 or BDNF (2^nd^ factors) and TRK/FYN coupling was analyzed. Interestingly, two-way ANOVA indicated a significant interaction between ANG2 and PD123319 (F_1,19_=5.08, p<0.05) and, surprisingly, between BDNF and PD123319 (F_1,20_=6.74, p<0.05). Respectively, pairwise comparisons showed PD123319 abolished ANG2 and BDNF effect of increasing TRK/FYN coupling (Fisher’s LSD, NS for both; Figure 4a). Next, we measured if PD123319 or ANG2 was able to modulate surface levels of TRKB. In fact, cultured cortical cells exposed to PD123319 presented decreased, while ANG2 increased, surface levels of TRKB (F_2,30_=24.33; Figure 4c).

**Figure 4.**
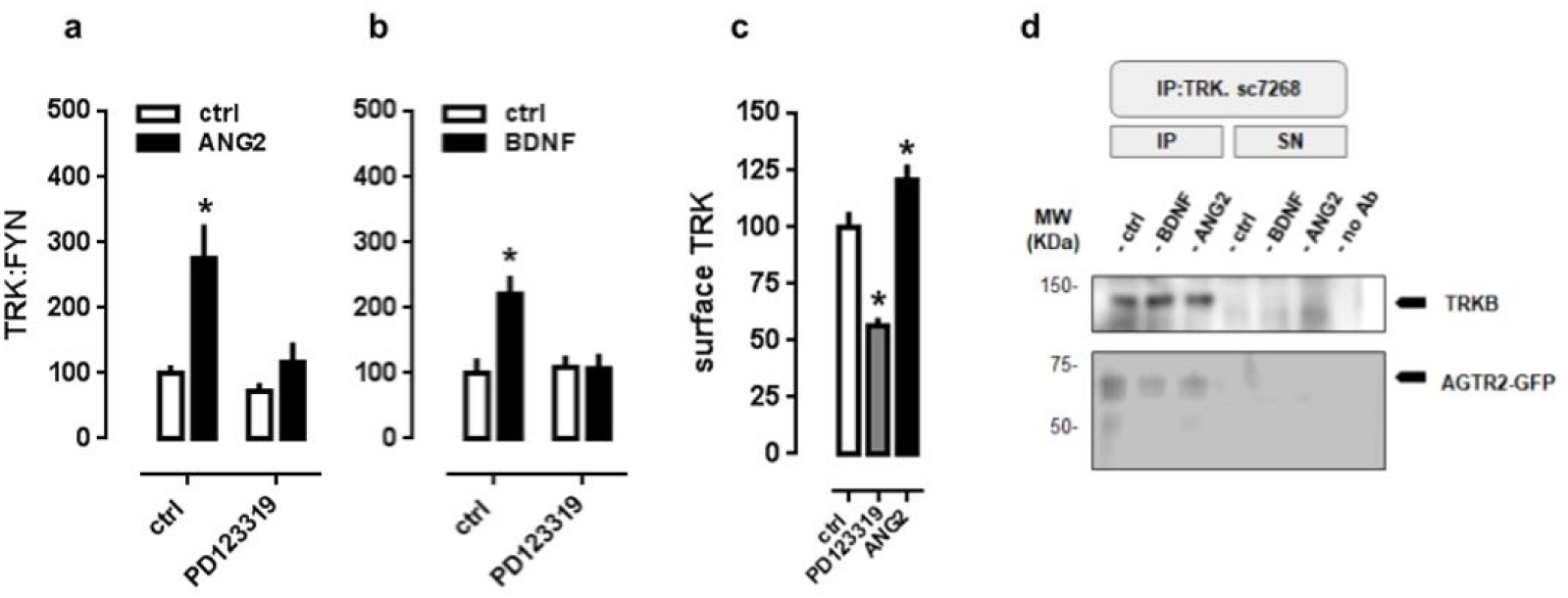
AGTR2-dependent interaction with FYN, surface positioning of TRKB and co-immunoprecipitation of AGTR2 and TRKB. Previous treament with PD123319 impaired TRK:FYN coupling induced by (a) ANG2 (n=5-6/group) or by (b) BDNF (n=6/group). (c) PD123319 decreased while ANG2 increased the levels of TRKB in the surface of cultured cortical cells (n=9-12/group). (d) Sample from MG87.TRKB fibroblast cell line, overexpressing AGTR2 labelling on blotting membrane from immunoprecipitation of TRKB protein. *p<0.05 compared to ctrl/ctrl group, unless otherwise stated; cells were lysed 10min after last drug administration.

Altogether, these results point to the existence of a heterodimer TRKB/AGTR2, since previous studies have described cross-antagonism (ability of both antagonists of each receptor units in the heterodimer to block signaling of each other agonist) as a fingerprint of heterodimerization (Ellis et al., 2006; Ferrada et al., 2009). In agreement with this idea, as observed in Figure 4d, a labeled GFP-tagged AGTR2 was co-precipitated with TRKB, however no apparent effect of BDNF or ANG2 was found in the levels of such complex.

### Challenge with glutamate inverts the ratio between AGTRs in cortical cells

Preliminary comparison between the presented *in vitro* and *in vivo* analysis indicates an incompatibility regarding the effects of losartan. This compound, although effective when injected systemic or into mPFC, did not exerted any effect *per se* in cultured cortical cells. Therefore, we considered the putative role of pretest stress in animals. In this scenario, a single exposure to inescapable stress, in addition to a peak in corticosterone production (lasting for 2h), increases glutamate levels for up to 24h (Musazzi et al., 2016; Popoli et al., 2011). Therefore, firstly we determined the levels of AGTR1 and AGTR2 mRNA in our cultured cells. The results observed in figure 5a, indicated a 5-times higher expression of AGTR2 compared to AGTR1 (Mean ΔCt value/SEM(n); AGTR1: 17.08/0.09(3); AGTR2: 14.69/0.37(3); t_4_= 2.96, p<0.05). Further, we incubated cortical cells with glutamate (10 or 100uM/2h) and determined the levels of AGTRs mRNA. Separate analysis of AGTRs expression following glutamate insult suggests a decrease in AGTR2 mRNA levels (Mean of fold change/SEM from ctrl: 1.00/0.09; glutamate 10uM: 0.88/0.06 and glutamate 100uM: 0.47/0.10; n=5,5,4 respectively) but no change in AGTR1 (fold change from ctrl: 1.00/0.10; glutamate 10uM: 1.39/0.14 and glutamate 100uM: 1.12/0.11; n=5,5,4 respectively). In this condition, there was an inversion in the ratio between AGTR1 and AGTR2 mRNA levels [glutamate 10uM: 1.5-times more AGTR1; glutamate 100uM: 2.54-times more AGTR1], as seen in figure 5b. Then, cortical cells pre-exposed to glutamate (100uM/2h) were tested 24h after the insult for the interaction between losartan and ANG2. Interestingly, 24h after the insult with glutamate, putatively inverting the AGTR1/AGTR2 ratio, losartan was necessary for the effect of ANG2 on pTRK levels (interaction: F_1,20_= 14.46 p<0.05, Figure 5c).

**Figure 5.**
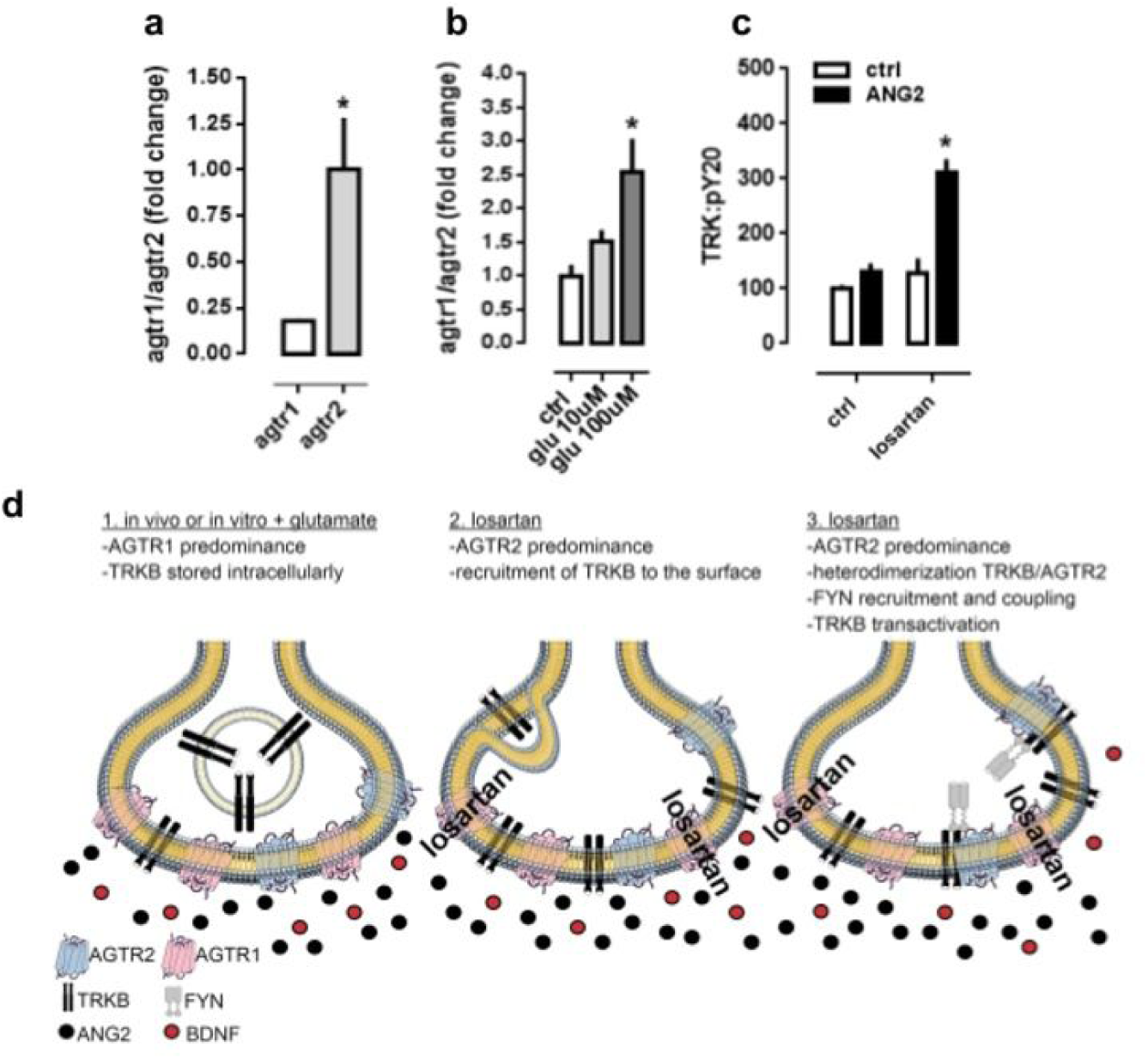
Challenge with glutamate (10 or 100uM/2h) inverts the levels of AGTRs in cortical cells. (a) AGTR2 subtype has a 5-times higher mRNA expression than AGTR1 (n=3/group). (b) The incubation with glutamate invert this ratio in favor of AGTR1. (c) Cells challenged with glutamate (100uM/2h) and tested for ANG2-induced activation of TRKB 24h later were responsive only in the presence of losartan (n=6/group). *p<0.05 from ctrl or agtr1 group. (d) Graphical abstract for the TRKB-dependent antidepressant-like effect of losartan. In basal conditions the low levels of AGTR1 compared to AGTR2 in cortical cells would be responsible to keep TRKB at the cell surface, passive to be activated by BDNF or transactivated by AGTR2. Upon excessive glutamatergic firing under stressful situations, the decrease in AGTR2 levels compromises TRKB activation. treament with losartan, putatively blocking AGTR1, favors the activation of AGTR2, reinstating TRKB activity.

## DISCUSSION

The present study indicates that systemic treatment with angiotensin receptor blocker (ARB) losartan and imipramine (used as positive control), but not with angiotensin converting enzyme inhibitors (ACEi) captopril or enalapril, promoted antidepressant-like effect in the FST. Losartan infusion into vHC, or into PL, but not into dHC, was enough to mimic antidepressant-like effect of systemic injection. Both hippocampus and vmPFC are core structures to modulate motivational and emotional behavioral consequences of stress exposure, including depressive disorders (Castrén, 2005; Nestler et al., 2002). In agreement with our data, vHC is suggested to be mainly related to behavioral and physiological consequences of stress exposure, while dHC engages cognitive and learning process concerning spatial navigation (Fanselow and Dong, 2010). The antidepressant-like effect of losartan probably rely on TRK signaling acting in hippocampus and vmPFC prelimbic aspects (PL), whereas AGTR2 activation is required only in PL, since TRK inhibitor into vHC and PL, and AGTR2 antagonist into PL decreased that effect. Moreover, FST data could be a misleading from locomotor activity, since no change was observed in this parameter following the pharmacological treatments (Figure S1).

The next behavioral approach used in this study reinforces the involvement of BDNF/TRKB system in the effect of losartan. In mice, with dampened BDNF expression, losartan was no longer able to exert antidepressant-like effects. Similarly to the observed for losartan, these mice are not responsive to classical antidepressants, such as imipramine and fluoxetine (Castrén and Antila, 2017; Karpova et al., 2011; Saarelainen et al., 2003).

In order to corroborate behavioral experiments and further explore the mechanisms involved in losartan effects, we used primary cultures of embryonic cortex to evaluate interaction between RAS and BDNF/TRKB signaling. First, we observed that K252a and PD123319, but not losartan, prevented ANG2 effect of increasing pTRK levels, thus suggesting that ANG2 increases pTRK levels putatively by acting on AGTR2. This possibility is strengthened by the effect of AGTR2 agonist CGP42112 also inducing an increase in TRK activation, which is counterbalanced by previous administration of AGTR2 antagonist PD123319. Since the soluble form of TRKB (TRKB.Fc) did not prevent ANG2 effect on pTRK, it is plausible to consider no increment in BDNF release in this process. In this sense, transactivation of TRKB, or a facilitatory effect of basal levels of BDNF on its receptor, are reasonable scenarios. In fact, previous studies have described that both GPCR ligands adenosine and pituitary adenylate cyclase-activating polypeptide can transactivate TRK (Rajagopal et al., 2004). In addition, TRK transactivation by adenosine 2A receptor agonist was blocked by PP1, suggesting an involvement of SRC family tyrosine kinase (Lee and Chao, 2001). Later, FYN was described as the SRC member responsible for TRK transactivation (Huang and McNamara, 2010; Rajagopal and Chao, 2006). Accordingly, lipid raft localization of TRKB is regulated by FYN (Pereira and Chao, 2007). In agreement with this evidence, we observed ANG2 was also able to increase levels of TRK/FYN coupling in cortical cultures. Therefore, we propose that FYN acts as an intermediary molecule capable of inducing TRKB transactivation when ANG2 acts on AGTR2. Moreover, it was described that BDNF itself could promote a greater TRKB/FYN coupling (Iwasaki et al., 1998). Corroborating that prospect, our data also showed BDNF increasing TRKB/FYN coupling. Therefore, both ANG2 or BDNF, which are able to increase pTRK levels, also induce TRK/FYN coupling.

In addition, as expected PD123319 blocked TRKB/FYN coupling from ANG2 action, but unexpectedly PD123319 also prevented such coupling from BDNF action. Unexpected as well was the data indicating that PD123319 prevents BDNF effect of increasing pTRKB levels. Besides, generalized interaction of AGTR2 with other TRK members is unlikely, given that PD123319 did not prevent NFG action of increasing pTRKA levels. These unforeseen interactions can be explained by the observation that PD123319 is able to reduce surface expression of TRKB, whereas ANG2 leads to an increase, thereby suggesting a putative displacement of TRK to surface upon AGTR2 signaling and the decrease of BDNF effectiveness with previous PD123319. Indeed, modulation of TRK surface trafficking is important considering the two possible scenarios described above for the activation of TRKB, which might happen on cell membrane (Rajagopal et al., 2004). In addition, MG87.TRKB cell line, which overexpresses TRKB, allowed us to observe co-immunoprecipitation of GFP-tagged AGTR2 and TRKB, suggesting an AGTR2/TRKB dimerization. This approach was chosen for two main reasons: as analyzed by the group of Juan Saavedra, commercially available antibodies against AGTRs are far from ideal (Hafko et al., 2013); second, the cell line used expresses exclusively TRKB, thus being an ideal tool to our purpose.

Preliminary analysis showed that *agtr2* mRNA levels is 5 times higher than *agtr1a* in our primary cultures, and this ratio is inverted to 2.5 times more *agtr1a* after an insult with glutamate, and this later feature seems to allow a cooperative effect of losartan and ANG2. Using a model of retinal ischemia, it was observed increased expression of *agtr1a* mRNA peaked 12h after reperfusion, while the treatment with candesartan was able to prevent ischemia-induced glutamate release (Fujita et al., 2012). Taken together, these data indicate a possible positive feedback between AGTR1 signaling and glutamatergic transmission. Moreover, in line with our *in vitro* observations, the levels of *agtr1a* was increased while *agtr2* was decreased in medulla of stress-induced hypertensive rats (Du et al., 2013). However, an opposed effect of glutamate on *agtr2* mRNA have also been described (Makino et al., 1998). In this study, the insult with glutamate led to an increase in *agtr2* mRNA. The precise mechanism where stressful events or excessive glutamate release might reduce the levels of *agtr2* are still not comprehended and these apparent discrepancies could rest on methodological differences. For example, the culture method of Makino and colleagues rely on cortical cells cultivated for 14 days, supplemented with calf serum and mitosis inhibitors; while our cultures were serum-free (substituted by B27), cultivated for 8 days without any drugs to prevent cell proliferation.

Given that the FST protocol for mice does not engage the exposure to a previous stressful session, another scenario poses that the levels of *agtr1a* are kept higher by basal glutamatergic or simply by basal activity of cortical area. To address this possibility we used the EMBL-EBI Expression Atlas (http://www.ebi.ac.uk/gxa/home) to compare the expression of *agtr1a* and *agtr2* in murine brain. We only obtained data from RNAseq experiments conducted in four strains of mice by Gregg and colleagues (Gregg et al., 2010). In line with the proposed scenario, these authors observed that the levels of *agtr2* are lower or at least equivalent to *agtr1a* in prefrontal cortex of adult mice. Our behavioral data, indicating the effectiveness of losartan in mice submitted to FST, also suggests that the *agtr1a/2* ratio *in vivo* favors *agtr1a*, since no previous stressful event is presented in this protocol.

In conclusion, our findings indicate that losartan induces antidepressant-like effect possibly mediated by AGTR2 and TRKB coupling in the mPFC. We disclose a previously unknown TRKB activation by AGTR2, involving recruitment of SRC family kinase FYN. According to our data, and as summarized by the graphical abstract in figure 5, TRKB and AGTR2 form a heterodimer that probably docks FYN kinase to promote a crosstalk, putatively inducing pTRKB/PLCγ1 signaling. Under stressful conditions or excessive glutamatergic transmission, the increase in AGTR1 levels prevents the ANG2-triggered docking of FYN to the AGTR2/TRKB complex. Losartan, by blocking AGTR1, restore the AGTR2-mediated effects of ANG2 on TRKB. Considering the high comorbidity between depression and cardiovascular disorders, our results suggest ARBs, such as losartan, could be a therapeutically interesting tool in these cases, or even be available as a novel class of antidepressant drugs.

## ACKNOWLEDGMENTS

We thank Flávia Salata (USP), Sulo Kolehmainen (UH) and Outi Nikkila (UH) for their technical support; and Dr. Wing-Tai Cheung (from The Chinese University of Hong Kong) by kindly providing the plasmid construct coding for GFP-tagged AGTR2 for transfection.

